# Tdrd3 regulates the progression of meiosis II through translational control of *Emi2* mRNA in mouse oocytes

**DOI:** 10.1101/2021.02.17.431574

**Authors:** Natsumi Takei, Keisuke Sato, Yuki Takada, Rajan Iyyappan, Andrej Susor, Takehiro Yamamoto, Tomoya Kotani

**Author notes:** Corresponding author. Tomoya Kotani, Department of Biological Sciences, Faculty of Science, Hokkaido University, North 10 West 8, Sapporo, Hokkaido 060-0810, Japan.

## Abstract

After completion of meiosis I, the oocyte immediately enters meiosis II and forms a metaphase II (MII) spindle without an interphase, which is fundamental for generating a haploid gamete. Here, we identify tudor domain-containing protein 3 (Tdrd3) as a novel regulator of oocyte meiosis. Although early mitotic inhibitor 2 (Emi2) protein has been shown to ensure the meiosis I to II transition and the subsequent MII spindle formation by inhibiting the anaphase-promoting complex/cyclosome (APC/C), how it accumulates after meiosis I has remained unresolved. We isolated Tdrd3 as a protein directly binding to *Emi2* mRNA. In GV-stage mouse oocytes, *Emi2* mRNA assembled into RNA granules containing Tdrd3, while cyclin B1 mRNA, which was translated in early meiosis I, formed different granules. Knockdown of Tdrd3 attenuated Emi2 synthesis in meiosis II without affecting cyclin B1 synthesis in meiosis I. Moreover, Tdrd3-deficient oocytes entered interphase and failed to form an MII spindle after completion of meiosis I. Taken together, our results indicate the importance of Tdrd3-mediated translational control of *Emi2* mRNA, which promotes Emi2 synthesis in meiosis II, for the progression of meiosis.

## INTRODUCTION

Gametes are generated through the two successive meiotic cell divisions, meiosis I and meiosis II. In oocytes, meiosis is arrested at the prophase of meiosis I, and more than ten thousand mRNAs are accumulated for oocyte growth and for the progression of meiosis and embryonic development (Conti and Franciosi, 2018; Masui and Clarke, 1979; Meneau et al., 2020; Winata and Korzh, 2018). Fully grown germinal vesicle (GV)-stage oocytes resume meiosis in response to hormonal stimulation. Shortly after the resumption of meiosis, oocytes activate maturation/M-phase-promoting factor (MPF), which consists of the catalytic subunit Cdc2 and regulatory subunit cyclin B (Nurse, 1990). The kinase activity of MPF promotes chromosome condensation, dissolution of the nuclear membrane, and formation of a metaphase I (MI) spindle (Abrieu et al., 2001; Kotani and Yamashita, 2002; Ledan et al., 2001; Polanski et al., 1998). Degradation of cyclin B by anaphase-promoting complex/cyclosome (APC/C) partially inactivates MPF, resulting in the separation of homologous chromosomes. Subsequently, the MPF activity is rapidly increased, and oocytes enter meiosis II and form a metaphase II (MII) spindle without an interphase. In vertebrates, meiosis is arrested again at MII by an activity called cytostatic factor (CSF) while awaiting fertilization (Masui and Markert, 1971; Sagata, 1996). CSF stabilizes MPF activity indirectly by inhibition of APC/C, preventing sister chromatid separation at MII (Madgwick et al., 2004; Nixon et al., 2002).

Early mitotic inhibitor 2 (Emi2) has been identified as one of the components of CSF in *Xenopus* and mouse oocytes (Madgwick et al., 2006; Schmidt et al., 2005; Shoji et al., 2006; Tung et al., 2005). Emi2 was shown to inhibit APC/C by binding directly to APC/C, resulting in the arrest of meiosis at MII in *Xenopus* (Ohe et al., 2010). Emi2 protein is absent in GV-stage immature *Xenopus* oocytes and accumulates in meiosis II (Liu et al., 2006; Ohe et al., 2007). Previous studies showed that inhibition of Emi2 synthesis by injection of *Emi2* small interfering RNA (siRNA) or antisense morpholino oligo nucleotide (MO) resulted in failure in arrest at MII, while ectopic expression of Emi2 in meiosis I resulted in the arrest of meiosis at MI in *Xenopus* and mouse oocytes (Madgwick et al., 2006; Ohe et al., 2007; Shoji et al., 2006; Suzuki et al., 2010). These studies indicate that the timing of Emi2 synthesis as well as Emi2 synthesis itself is critical for APC/C inhibition in meiosis II. However, how and when Emi2 is synthesized from the mRNA stored in oocytes remain unresolved.

Of the mRNAs that have accumulated in GV-stage oocytes, thousands of mRNAs have been shown to be translationally repressed after transcription (Chen et al., 2011). After the resumption of meiosis, hundreds of mRNAs begin to be translated in meiosis I and more than one thousand mRNAs begin to be translated in meiosis II (Chen et al., 2011; Luong et al., 2020). cyclin B1 mRNA is one of the mRNAs translated in early meiosis I (Haccard and Jessus, 2006; Han et al., 2017; Kotani et al., 2013; Nakahata et al., 2003; Yang et al., 2017). The 3’ untranslated region (3’UTR) of many dormant mRNAs including cyclin B1 mRNA contains the cytoplasmic polyadenylation element (CPE), which is recognized by CPE-binding proteins (CPEBs) (Richter, 2007). CPEB1 has been shown to mediate the cytoplasmic polyadenylation of dormant mRNAs with short poly(A) tails after resumption of meiosis, resulting in translational activation of target mRNAs (Gebauer et al., 1994; Hake and Richter, 1994; Stebbins-Boaz et al., 1996; Tay et al., 2000). Although a large number of dormant mRNAs contain CPEs in their 3’UTRs, they are translated in different periods, suggesting that other mechanisms in addition to the mechanism that depends on the CPEB1 regulate the timing of translational activation of distinct mRNAs.

Recent studies have shwon that cytoplasmic RNA granules such as stress granules and processing bodies (P-bodies) mediate post-transcriptional regulations including stabilization and degradation of assembled mRNAs (Anderson and Kedersha, 2009; Buchan and Parker, 2009). We previously showed that the translation of cyclin B1 mRNA in meiosis I was regulated by the formation and disassembly of RNA granules (Kotani et al., 2017; Kotani et al., 2013). *Mad2* mRNA was shown to be translated in a similar period, and both mRNAs were targets of an RNA-binding protein, Pumilio1 (Pum1) (Takei et al., 2020). Pum1 was shown to form aggregates in GV-stage oocytes and regulate the translation of cyclin B1 and *Mad2* mRNAs through assembly and dissolution of the aggregates.

The 3’UTR of *Xenopus emi2* mRNAs contains several CPEs that were shown to bind to CPEB1 (Tung et al., 2007). Although CPEB1 was shown to direct the polyadenylation of many dormant mRNAs after resumption of meiosis, this protein is degraded in meiosis I in *Xenopus* and mouse oocytes (Chen et al., 2011; Mendez et al., 2002; Setoyama et al., 2007). In *Xenopus* oocytes, polyadenylation of dormant mRNAs in meiosis II was shown to be directed by CPEB4, which accumulated in the later stage of meiosis I (Igea and Mendez, 2010). However, injection of antisense *Cpeb4* MO into GV-stage oocytes had no significant effect on the progression of meiosis in mouse oocytes (Chen et al., 2011), suggesting that CPEB4 is not mainly involved in the *Emi2* mRNA translation in mouse oocytes.

In this study, we identified tudor domain-containing protein 3 (Tdrd3) as a novel protein that binds to *Emi2* mRNA in mouse oocytes. *Emi2* mRNA was found to assemble into granules, which were different from those of cyclin B1 mRNA. Although HuR and HuB, components of stress granules (Gallouzi et al., 2000; Markmiller et al., 2018), and CPEB1 interacted with both *Emi2* and cyclin B1 mRNAs, Tdrd3 bound to and colocalized with *Emi2*, but not cyclin B1, mRNA. Knockdown of Tdrd3 attenuated Emi2 synthesis in meiosis II but did not affect the synthesis of cyclin B1 and Mad2 in meiosis I. Consistent with this, Tdrd3-deficient oocytes entered interphase and failed to form an MII spindle after completion of meiosis I. Taken together, our results indicate that the assembly of Tdrd3-containing granules controls the translation of *Emi2* mRNA in meiosis II, through which oocytes promote the progression of meiosis II.

## RESULTS

### Temporally controlled translation of *Emi2* mRNA is prerequisite for the generation of a fertilizable gamete

To assess how the function of Emi2 is temporally regulated in meiosis, we first analyzed the accumulation of Emi2 protein in mouse oocytes by raising antibodies against the N-terminal region of mouse Emi2 (amino acids 1-366). Affinity-purified anti-Emi2 antibodies recognized a protein with an apparent molecular mass of ~90 kDa in immunoblots of MII-stage oocytes but not GV-stage oocytes (Fig. S1A), which corresponds to the size of Emi2 shown in a previous study (Madgwick et al., 2006). The intensity of this signal was significantly reduced when the antibody was preincubated with recombinant Emi2 (Fig. S1B) and when the translation of *Emi2* mRNA was prevented by the antisense MO (Fig. 1D), confirming that the antibodies specifically recognize Emi2. Emi2 was not detected at 0 h (GV stage) and 8 h (MI stage) after resumption of meiosis, while a large amount of Emi2 was detected at 18 h (MII stage) (Fig. 1A). In contrast, the amount of cyclin B1 was increased at the MI stage and peaked at the MII stage (Fig. 1A), consistent with the results in previous studies (Ganesh et al., 2020; Takei et al., 2020; Yang et al., 2017).

**Fig 1.**
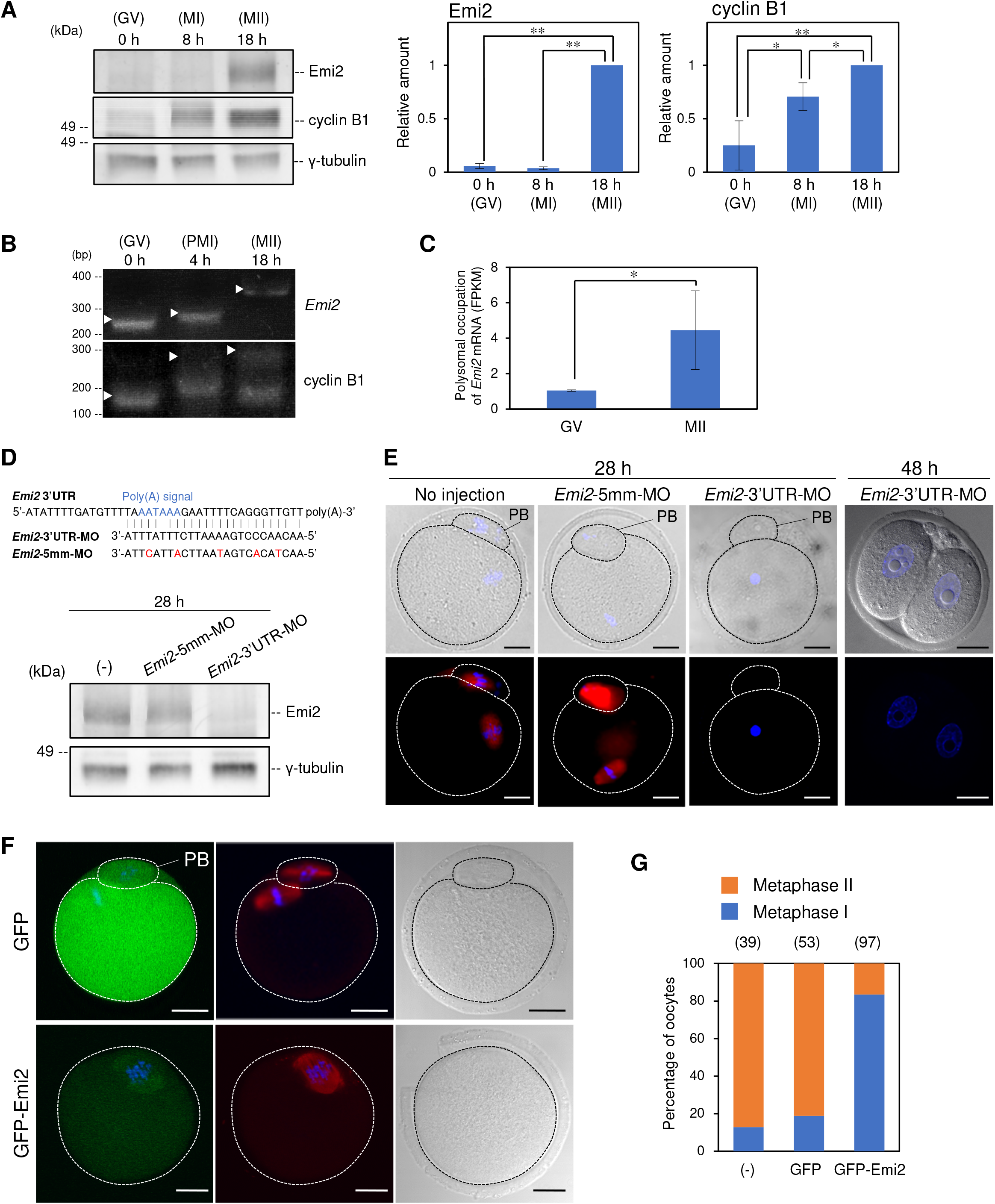
Polyadenylation-dependent translational control of *Emi2* mRNA defines the timing of Emi2 accumulation in mouse oocytes. (A, left) Immunoblotting of Emi2, cyclin B1 and γ-tubulin in oocytes at 0, 8 and 18 h after resumption of meiosis. (right) Quantitative analysis (means ± SD; n = 3). The statistical significance was analyzed by the Tukey-Kramer test. * *P* < 0.05, \*\**P* < 0.01. (B) Time course of the PAT assay of *Emi2* (upper) and cyclin B1 (lower) mRNAs after resumption of meiosis. Similar results were obtained from three independent experiments. Note that poly(A) tails of a fraction of cyclin B1 mRNA were shortened in MII oocytes, possibly due to the induction of meiosis *in vitro*. (C) RNA-seq-based gene expression values (FPKM) for *Emi2* in GV- and MII-stage oocytes (means ± SD; n = 3). *t*-test: * *P* < 0.05. (D, upper) Sequences of the 3’UTR of *Emi2* mRNA that are targeted by *Emi2*-3’UTR-MO. *Emi2*-5mm-MO contains 5 mismatches (red). (D, lower) Immunoblotting of Emi2 and γ-tubulin in oocytes not injected (-) and injected with *Emi2*-5mm-MO and *Emi2*-3’UTR-MO at 28 h after resumption of meiosis. (E) Immunofluorescence of β-tubulin (red) in oocytes injected with *Emi2*-5mm-MO and *Emi2*-3’UTR-MO at 28 h and 48 after resumption of meiosis. DNA is shown in blue. (F) Immunofluorescence of β-tubulin (red) in oocytes injected with GFP or GFP-Emi2 mRNA at 18 h after resumption of meiosis. Right figure shows a bright field. DNA is shown in blue. (G) Percentage of metaphase I- or metaphase II-stage oocytes at 18 h after resumption of meiosis. The numbers in parentheses indicate the total numbers of oocytes analyzed. PB, polar body. Bars: 20 μm.

A poly(A) test (PAT) assay showed that *Emi2* mRNA was polyadenylated only slightly at 4 h (prometaphase I, PMI, stage) and was significantly polyadenylated at 18 h (MII stage) (Fig. 1B). In contrast, cyclin B1 mRNA was rapidly polyadenylated at the PMI stage and polyadenylation peaked at the MII stage (Fig. 1B). Polysomal fractionation analysis showed that the amount of *Emi2* mRNA in the polysomal fraction increased by approximately 5 fold from the GV stage to the MII stage (Fig. 1C), although monosomal and polysomal occupation of *Emi2* mRNA was equal between the GV and MII stages (Fig. S1C). Taken together, these results indicate that *Emi2* mRNA is translationally repressed in meiosis I with short poly(A) tails and is translated in meiosis II possibly through the polyadenylation of mRNA.

A recent study demonstrated that antisense MOs targeting the 3’-end sequences of mRNAs prevent the polyadenylation of mRNAs (Wada et al., 2012). To assess the importance of polyadenylation of *Emi2* mRNA, we injected an antisense *Emi2* MO that targets the terminal sequence of the *Emi2* mRNA 3’UTR (*Emi2*-3’UTR-MO) and an *Emi2* MO that contains 5-nts mismatches (*Emi2*-5mm-MO) as a control (Fig. 1D) into GV-stage oocytes. Immunoblot analysis showed that the synthesis of Emi2 was effectively inhibited by *Emi2*-3’UTR-MO, but not by *Emi2*-5mm-MO, at 28 h after resumption of miosis (Fig. 1D). Immunostaining showed that the control oocytes had extruded the first polar body and had formed an MII spindle with chromosomes aligned on the equatorial plate at 28 h after resumption of meiosis (Fig. 1E), indicating that the oocytes are arrested at MII. In contrast, the oocytes injected with *Emi2*-3’UTR-MO possessed a pronucleus-like structure with the first polar body (Fig. 1E). At 48 h after resumption of meiosis, the *Emi2*-3’UTR-MO-injected oocytes entered into the two-cell stage. These phenotypes were consistent with the defects observed in oocytes in which Emi2 was knocked down by *Emi2*-siRNA or *Emi2*-MO targeting the translational start site of *Emi2* mRNA (Madgwick et al., 2006; Shoji et al., 2006). These results indicate that Emi2 is translated in meiosis II in a polyadenylation-dependent manner, which is required for the entry into meiosis II and the prevention of parthenogenetic development.

We then confirmed the effect of precocious Emi2 synthesis in meiosis I by injecting mRNA encoding GFP or GFP-Emi2 into GV-stage oocytes. In oocytes expressing GFP, 81.1% of the oocytes underwent normal meiotic progression and were arrested at MII (Fig. 1F and G). In contrast, 83.5% of the oocytes expressing GFP-Emi2 were arrested at MI (Fig. 1F and G). In these oocytes, GFP-Emi2 was localized at the MI spindle (Fig.1F). These results are consistent with results of previous studies (Madgwick et al., 2006; Suzuki et al., 2010). Taken together, our results indicate that the temporally controlled translation of *Emi2* mRNA defines the timing and duration of Emi2 function, which is prerequisite for meiosis I to II transition, progression of meiosis II and arrest at MII.

### *Emi2* mRNA forms cytoplasmic granules that are distinct from cyclin B1 RNA granules

To elucidate the mechanisms by which the translation of *Emi2* mRNA is temporally regulated, we first analyzed the distribution of *Emi2* mRNA in mouse oocytes. Previous studies demonstrated that dormant cyclin B1 and *Mad2* mRNAs assemble into granules in GV-stage oocytes, and the disassembly of these granules after resumption of meiosis regulates the timing of translational activation of assembled mRNAs (Kotani et al., 2013; Takei et al., 2020). *Emi2* mRNA was detected in GV-stage oocytes by the highly sensitive *in situ* hybridization with the tyramide signal amplification (TSA) system (Fig. 2A). Fluorescent *in situ* hybridization (FISH) with the TSA system showed that *Emi2* mRNA formed RNA granules in the oocyte cytoplasm (Fig. 2B). Interestingly, double FISH analysis showed that *Emi2* and cyclin B1 mRNAs formed different granules (Fig. 2C).

**Fig 2.**
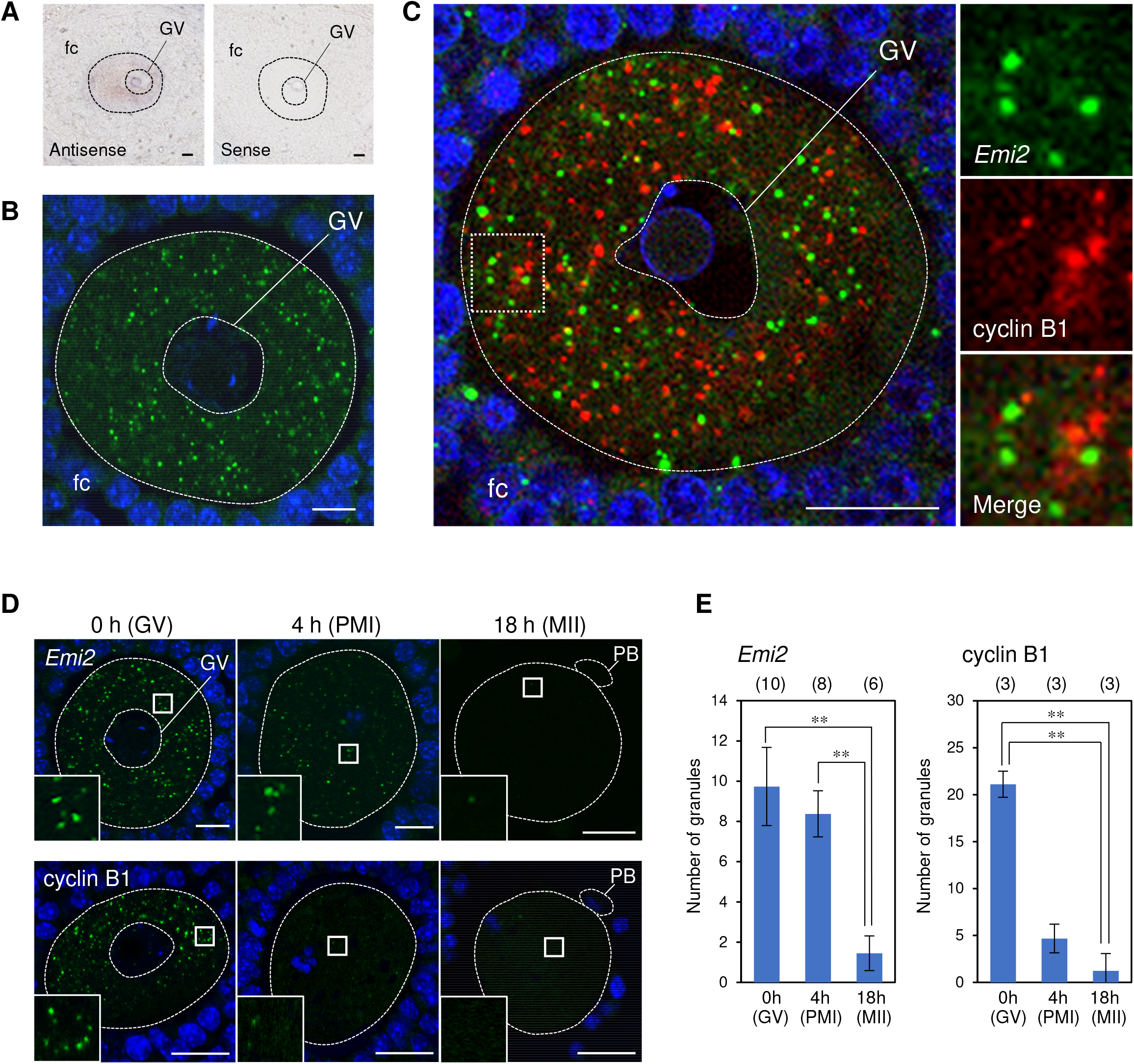
*Emi2* mRNA forms granules that are different from cyclin B1 RNA granules. (A) Expression of *Emi2* mRNA in the mouse ovary. A mouse ovary section hybridized with the *Emi2* antisense probe (left) and sense probe (right). (B) FISH analysis of *Emi2* mRNA in GV-stage oocytes. (C) FISH analysis of *Emi2* (green) and cyclin B1 (red) mRNAs in GV-stage oocytes. (insets) Enlarged views of the boxed region. (D) FISH analysis of oocytes at 0, 4, 18 h after resumption of meiosis. (insets) Enlarged views of the boxed region. (E) The numbers of RNA granules per 100 μm^2^ in individual oocytes at 0, 4, and 18 h were counted (mean ± SD). The numbers in parentheses indicate the total numbers of oocytes analyzed. The statistical significance was analyzed by the Tukey-Kramer test. \*\**P* < 0.01. fc, follicle cells; GV, germinal vesicle; PB, polar body. Bars, 10 μm in A, B and D, 20 μm in C.

After the resumption of meiosis, large parts of cyclin B1 RNA granules rapidly disassembled at PMI as observed in previous studies (Kotani et al., 2013; Takei et al., 2020), while the number of *Emi2* RNA granules was maintained at this stage (Fig. 2D and E). In the MII stage, *Emi2* RNA granules had almost completely disappeared as in the case of cyclin B1 (Fig. 2D and E). Since the total amount of *Emi2* mRNA was not decreased in MII-stage oocytes (Fig. S1C), the disappearance of *Emi2* RNA granules indicates granule disassembly. The disassembly of *Emi2* RNA granules coincided with the polyadenylation of *Emi2* mRNA (Fig. 1B). These results suggest that translation of *Emi2* mRNA is regulated through formation and disassembly of RNA granules as in the case of cyclin B1 and*Mad2* mRNA (Kotani et al., 2013; Takei et al., 2020). However, *Emi2* RNA granules disassembled in later stages compared to the rapid disassembly of cyclin B1 RNA granules, suggesting differences in the assembly and regulation of these granules.

### Pum1 binds to cyclin B1 mRNA, but not to *Emi2* mRNA, while HuR and HuB bind to both mRNAs

To address the difference and similarity of *Emi2* and cyclin B1 RNA granules, we analyzed RNA-binding proteins interacting with *Emi2* and/or cyclin B1 mRNAs. The previous study demonstrated that Pum1 interacts with cyclin B1 and *Mad2* mRNAs and regulates the translational activation of these mRNAs in early meiosis I (Takei et al., 2020). To examine whether Pum1 interacts with *Emi2* mRNA, we performed an immunoprecipitation assay followed by RT-PCR (IP/RT-PCR). As expected, cyclin B1 mRNA was detected in precipitation with an anti-Pum1 antibody, while neither *Emi2* mRNA nor *α-tubulin* mRNA was detected in the precipitation (Fig. 3A), indicating that *Emi2* mRNA is not a target of Pum1. Recent studies have demonstrated that different types of cytoplasmic granules share several components (see a review Buchan and Parker, 2009). We then analyzed the interaction of HuR and HuB proteins with *Emi2* and cyclin B1 mRNAs, since HuR and HuB are components of stress granules and are expressed in the ovary (Colombrita et al., 2013; Hinman and Lou, 2008) and HuR was shown to be colocalized with cyclin B1 RNA granules in zebrafish oocytes (Kotani et al., 2013). *Emi2* and cyclin B1 mRNAs, but not *α-tubulin* mRNA, were detected in precipitations with an anti-HuR or anti-HuB antibody (Fig. 3B and C), indicating that HuR and HuB bind to both *Emi2* and cyclin B1 mRNAs. These results suggest that HuR and HuB are components common to *Emi2* and cyclin B1 RNA granules, whereas Pum1 is specific to cyclin B1 RNA granules.

**Fig 3.**
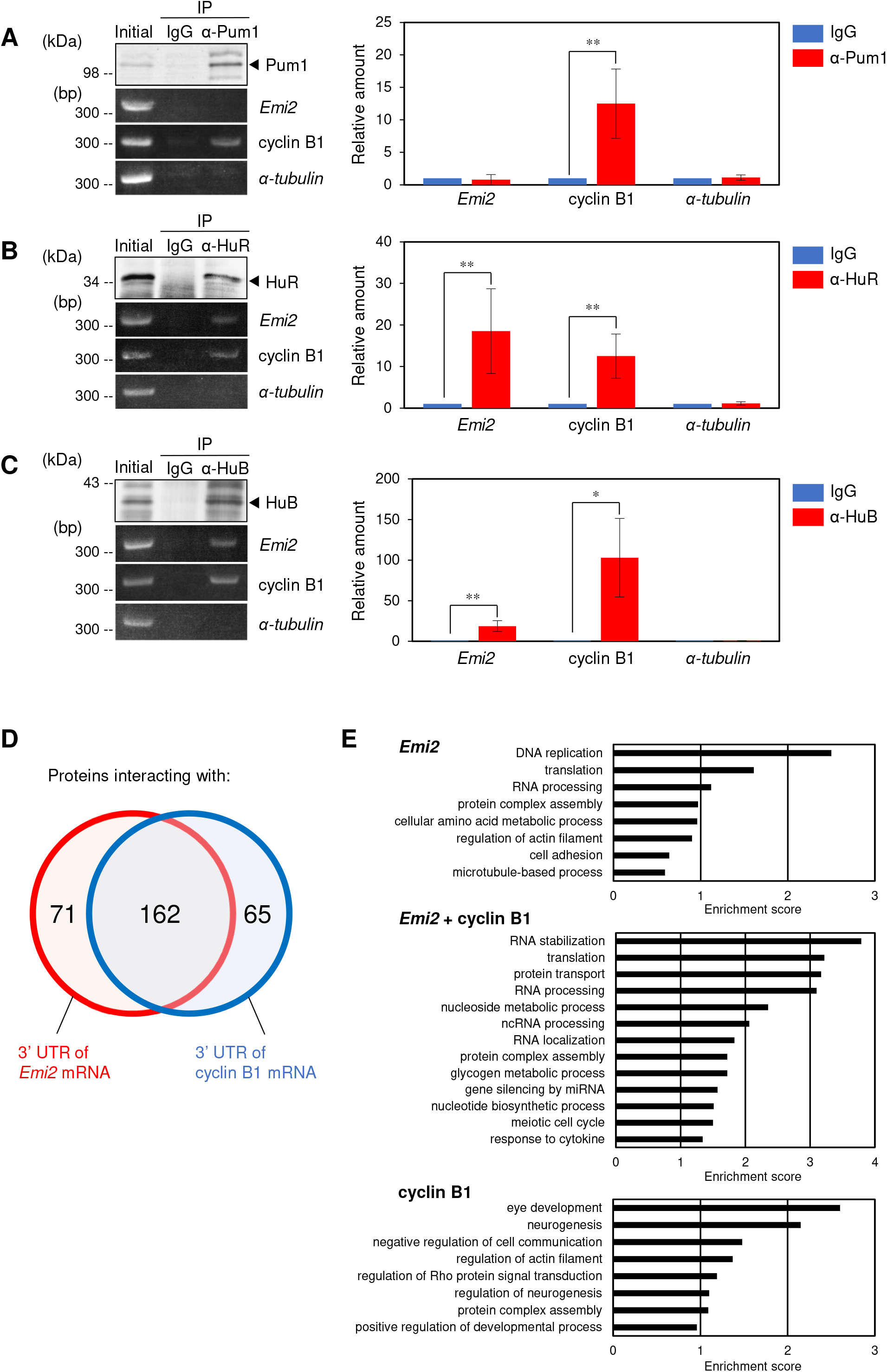
Specific and common interactions of RNA-binding proteins with *Emi2* mRNA. (A-C) Immunoblotting of mouse ovary extracts before IP (Initial) and IP with control IgG (IgG) or anti-Pum1 (α-Pum1) (A), anti-HuR (α-HuR) (B) and anti-HuB (α-HuB) (C) antibodies and semi-quantitative RT-PCR amplification for *Emi2*, cyclin B1 *and α-tubulin* transcripts. Graphs show the results of quantitative analyses of RT-PCR (means ± SD; n = 3). *t*-test: *\*P* < 0.05, *\*\*P* < 0.01. (D) Venn diagram depicts the number of proteins isolated as proteins interacting with 3’UTR sequences of *Emi2* (red) and cyclin B1 (blue) with the number of overlapping proteins. Keratin and ribosomal proteins were excluded. (E) Gene ontology analysis of genes enriched in proteins isolated as proteins interacting with 3’UTR sequences of *Emi2* (upper), *Emi2* and cyclin B1 (middle) and cyclin B1 (lower).

### Isolation of Tdrd3 as a protein directly binding to *Emi2*, but not cyclin B1, mRNA

The results showing that Pum1 binds to cyclin B1, but not *Emi2*, mRNA suggested that the precise regulation of distinct RNA granules is achieved by components specific to each RNA granule. To isolate proteins specific to *Emi2* RNA granules, we performed an RNA pull-down assay by using *in vitro*-synthesized RNAs of *Emi2* and cyclin B1 3’UTRs with oocyte extracts. In this assay, we used *Xenopus* oocytes since sufficient materials were not obtained by using mouse oocytes. Mass spectrometry analysis identified 71 candidate proteins specifically binding to *Emi2* mRNA, 162 candidate proteins binding to both mRNAs, and 65 candidate proteins specifically binding to cyclin B1 mRNA (Fig. 3D and Table S1). Gene ontology (GO) terms of the proteins were associated with RNA metabolism and translational control (Fig. 3E). Moreover, HuB and CPEB1 were isolated as proteins binding to both mRNAs, being consistent with the results of *in vivo* analysis (Fig. 3C) and of *in vitro* UV cross-linking assay (see Fig. 4C) and evaluating the results of RNA pull-down assay.

**Fig 4.**
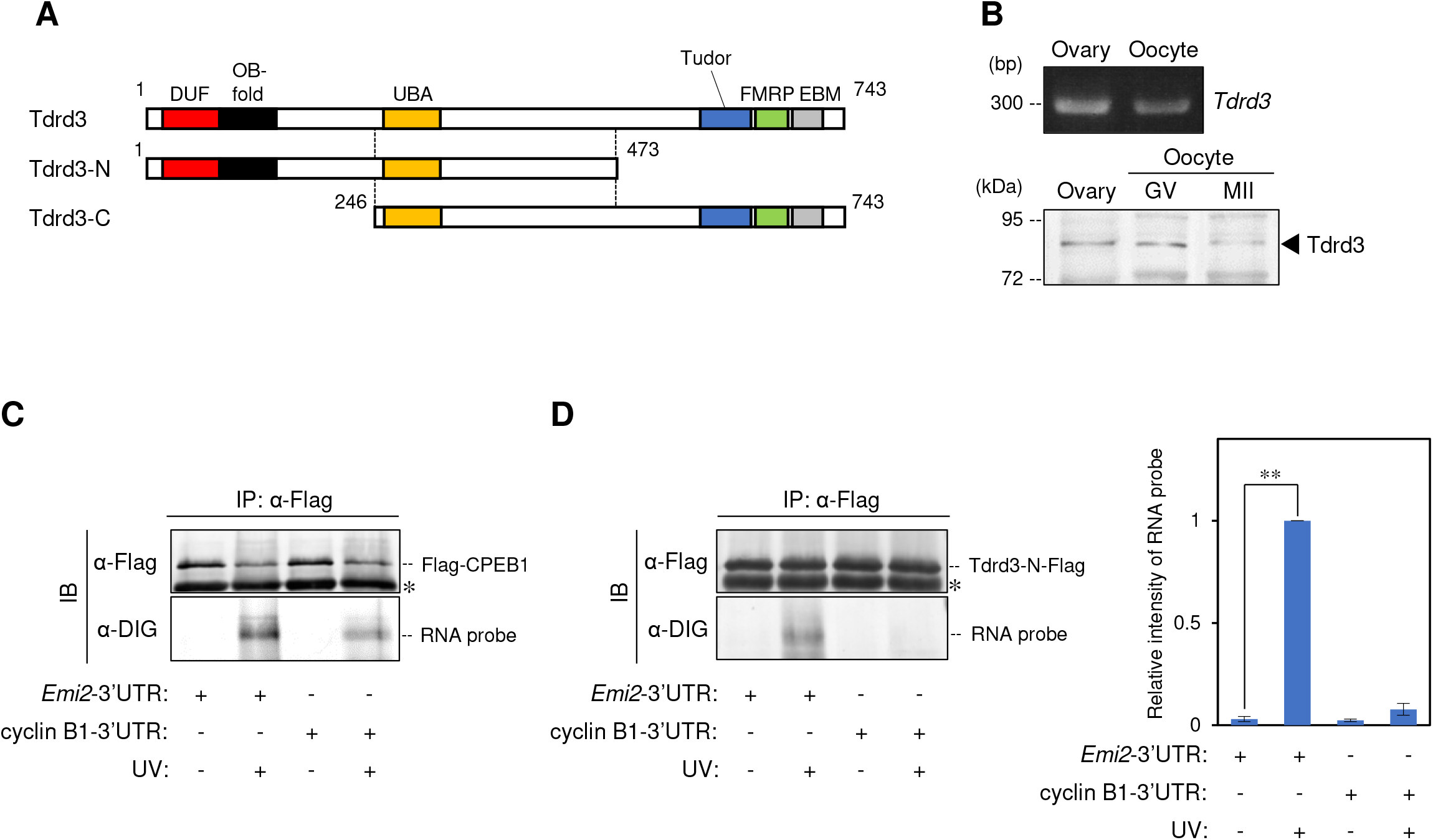
Tdrd3 is expressed in mouse oocytes and interacts with the 3’UTR of *Emi2* mRNA. (A) Schematic diagrams of Tdrd3, Tdrd3-N and Tdrd3-C. (B) RT-PCR amplification for *Tdrd3* mRNA in the ovary and oocytes and immunoblotting of Tdrd3 in extracts of the mouse ovary and oocytes at GV and MII stages. (C) UV-cross linking assay of the 3’UTR of *Emi2* and cyclin B1 mRNA with Flag-CPEB1. The upper panel shows immunoprecipitated Flag-CPEB1. The lower panel shows DIG-labeled probes that interact with Flag-CPEB1. An asterisk shows IgG of anti-Flag antibody. (D, left) UV-cross linking assay of the 3’UTR of *Emi2* and cyclin B1 mRNA with Tdrd3-N-Flag. The upper panel shows immunoprecipitated Tdrd3-N-Flag. The lower panel shows DIG-labeled probes that interact with Tdrd3-N-Flag. An asterisk shows IgG of anti-Flag antibody. (D, right) Quantitative analysis for the intensities of RNA probes (means ± SD; n =3).

Tudor domain-containing protein 3 (Tdrd3), a modular protein identified on the basis of its tudor domain of approximately 60 amino acids (Fig. 4A) (Goulet et al., 2008; Linder et al., 2008), was isolated as one of the candidates specifically binding to *Emi2* mRNA. Tdrd3 possesses putative nucleic acid recognition motifs (DUF and OB-fold motifs), a ubiquitin-binding associated (UBA) domain, a fragile X mental retardation protein (FMRP)-interacting domain, and an exon junction complex binding motif (EBM) (Fig. 4A). We focused on Tdrd3 since the expression of *Tdrd3* mRNA and that of Tdrd3 protein were confirmed in GV-stage mouse oocytes (Fig. 4B) and Tdrd3 has been shown to assemble into stress granules (Goulet et al., 2008; Linder et al., 2008), although its physiological function has remained unknown. Immunoblot analysis showed that the expression levels of Tdrd3 were not changed in GV- and MII-stage oocytes (Fig. 4B).

We first examined whether Tdrd3 directly binds to *Emi2* mRNA by performing an *in vitro* UV cross-linking assay, since a direct interaction between Tdrd3 and RNA has not been tested. RNA probes consisting of the 3’UTR of *Emi2* and cyclin B1 were incubated with Flag-tagged Tdrd3 and CPEB1. Both RNA probes were associated with CPEB1 (Fig. 4C), consistent with results of previous studies in *Xenopus* oocytes (Stebbins-Boaz et al., 1996; Tung et al., 2007). Since a large fraction of full-length Tdrd3 was destructed in this assay (Fig. S2A), we prepared deleted forms of Tdrd3 that consisted of N-terminal (1-473 aa) and C-terminal (246-743 aa) regions of Tdrd3 (Tdrd3-N and -C) (Fig. 4A). Immunoblot analysis showed that Tdrd3-C was divided into many fragments when it was translated in a rabbit reticulocyte lysate system regardless of the position of the Flag-tag, while Tdrd3-N was not destructed (Fig. S2B). Therefore, we used Flag-tagged Tdrd3-N for the UV cross-linking assay. The *Emi2* RNA probe was associated with Tdrd3-N-Flag, whereas the cyclin B1 RNA probe was not (Fig. 4D), indicating that Tdrd3 specifically and directly interacts with *Emi2* mRNA.

### Tdrd3 colocalized with *Emi2*, but not cyclin B1, RNA granules

To examine the interaction between Tdrd3 and *Emi2* mRNA *in vivo*, we first performed IP/RT-PCR by using anti-Tdrd3 antibodies. Unfortunately, Tdrd3 was not precipitated with antibodies raised against three different antigens. We then analyzed the distribution of Tdrd3 in the oocyte cytoplasm by immunostaining of mouse ovaries. Super-resolution microscopy (structured illumination microscopy, SIM) showed that Tdrd3 was distributed in the cytoplasm of GV-stage oocytes as many particles (Fig. 5A). Since these Tdrd3 particles seemed to form clusters in the cytoplasm, we calculated the average distance of five neighboring particles and compared it with that of randomly distributed dots (Fig. S3). The distance of Tdrd3 particles was significantly shorter than that of randomly distributed dots (Fig. 5C), suggesting the formation of Tdrd3 clusters. Simultaneous detections of Tdrd3 with Pum1 showed that particles of Tdrd3 were not overlapped with Pum1 aggregates (Fig. 5B). The size of Tdrd3 particles appeared to be smaller than that of Pum1 (Fig. 5D), and the shape of Tdrd3 particles tended to be round compared to that of Pum1 (Fig. 5E), which was shown to assemble into solid-like highly structured aggregates (Takei et al., 2020) (see also Fig. 5G). FISH analysis showed that Tdrd3 particles colocalized with *Emi2* RNA granules (Fig. 5F, arrows), but no overlap was observed with cyclin B1 RNA granules (Fig. 5F). In contrast, Pum1 aggregates surrounded and overlapped with cyclin B1 RNA granules (Fig. 5G, arrows), being consistent with the results of the previous study (Takei et al., 2020), while no overlap was observed with *Emi2* RNA granules (Fig. 5G), supporting the results of IP/RT-PCR analysis (Fig. 3A). These results demonstrate that Tdrd3 is a component specific to *Emi2* RNA granules.

**Fig 5.**
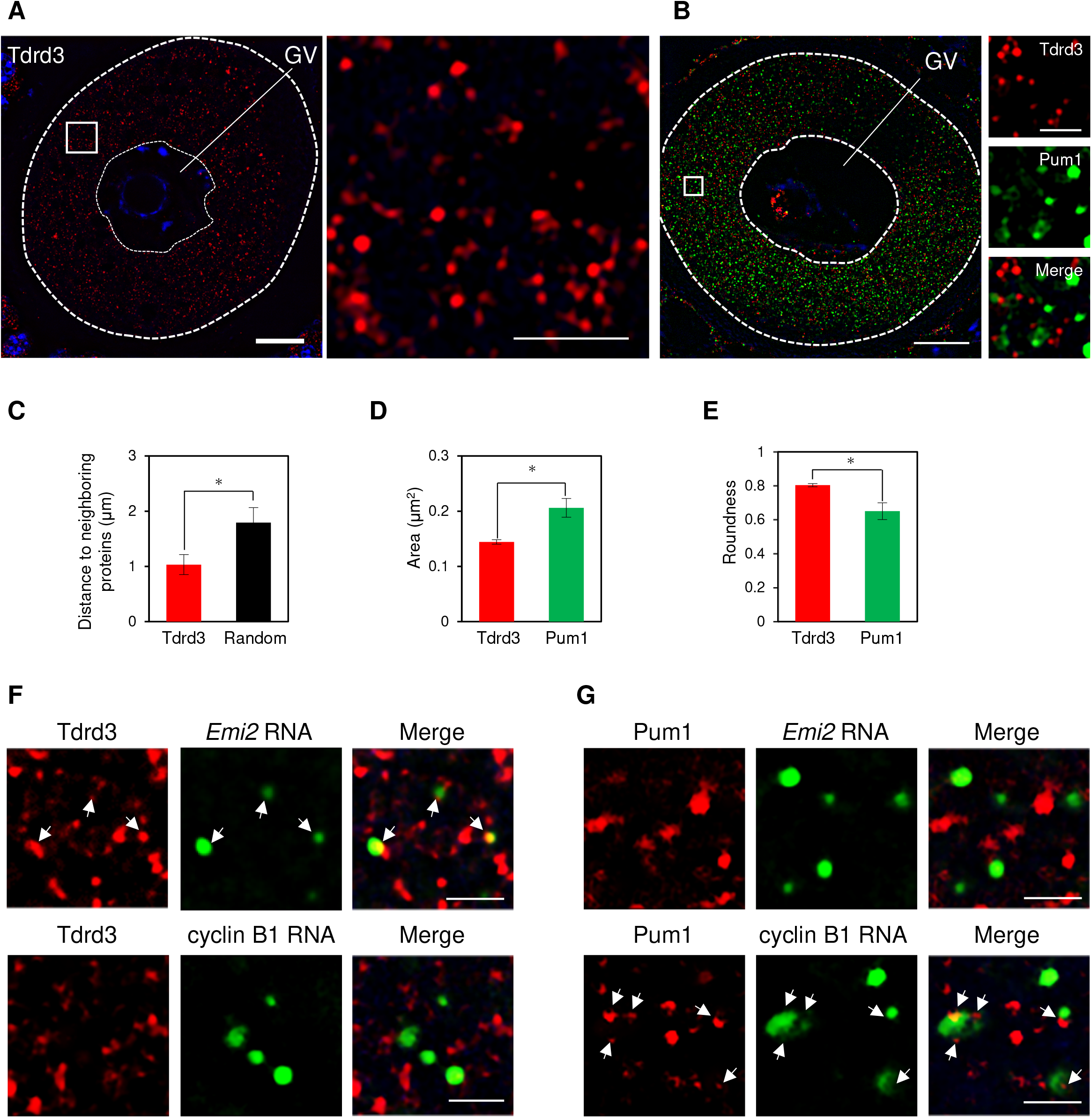
*Emi2* RNA granules contain Tdrd3. (A, left) Immunofluorescence of Tdrd3 in GV-stage oocytes. DNA is shown in blue. (right) An enlarged view of the boxed region. Similar results were obtained from three independent experiments. (B) Double immunofluorescence of Tdrd3 (red) and Pum1 (green) in GV-stage oocytes. (insets) Enlarged views of the boxed region. DNA is shown in blue. Similar results were obtained from three independent experiments. (C) Average distances (μm) between one Tdrd3 particle and 5 neighboring particles and average distances of random dots (n = 50 from three independent experiments). *t*-test: * *P* < 0.05. (D) Average sizes (μm^2^) of Tdrd3 and Pum1 (n = 90 from three independent experiments). *t*-test: * *P* < 0.05. (E) Roundness of Tdrd3 and Pum1 (n = 90 from three independent experiments). *t*-test: * *P* < 0.05. (F) Immunofluorescence of Tdrd3 (red) and FISH analysis of *Emi2* (upper, green) or cyclin B1 (lower, green) mRNAs. Arrows indicate Tdrd3 overlapped with *Emi2* RNA granules. (G) Immunofluorescence of Pum1 (red) and FISH analysis of *Emi2* (upper, green) or cyclin B1 (lower, green) mRNAs. Arrows indicate Pum1 overlapped with cyclin B1 RNA granules. GV, germinal vesicle. Bars: 10 μm in A and B, 2 μm in C and D.

### Tdrd3 controls the *Emi2* mRNA translation in meiosis II

To assess whether Tdrd3 plays a role in the regulation of *Emi2* mRNA, we injected an anti-Tdrd3 antibody or control IgG into GV-stage oocytes and induced the resumption of meiosis. Immunoblot analyses showed that the initial accumulation of Emi2 at 12 h after resumption of meiosis was prevented by the anti-Tdrd3 antibody (Fig. 6A and B), although the amount of Emi2 was recovered at 18 h. These results suggest that Tdrd3 is involved in the translational activation of *Emi2* mRNA in meiosis II. However, the oocytes injected with the anti-Tdrd3 antibody extruded a polar body in the normal time course and seemed to be arrested at MII as in the case of oocytes injected with IgG (Fig. S4A and B). We speculated that this antibody only partially interfered with the Tdrd3 function and that the recovery of Emi2 accumulation uncovered the actual function of Tdrd3 in the progression of meiosis. To overcome this, we first attempted to knock down Tdrd3 by antisense MOs, but injection of the antisense Tdrd3 MO did not reduce the amount of Tdrd3 even in oocytes incubated for 18 h after injection (Fig. S4C), suggesting that Tdrd3 is stable in oocytes. We then attempted to knock down Tdrd3 by using the Trim-Away protein degradation system (Clift et al., 2017) with the anti-Tdrd3 antibody. This system allows rapid depletion of an endogenous protein by TRIM-mediated degradation of the antibody-target protein complex.

**Fig 6.**
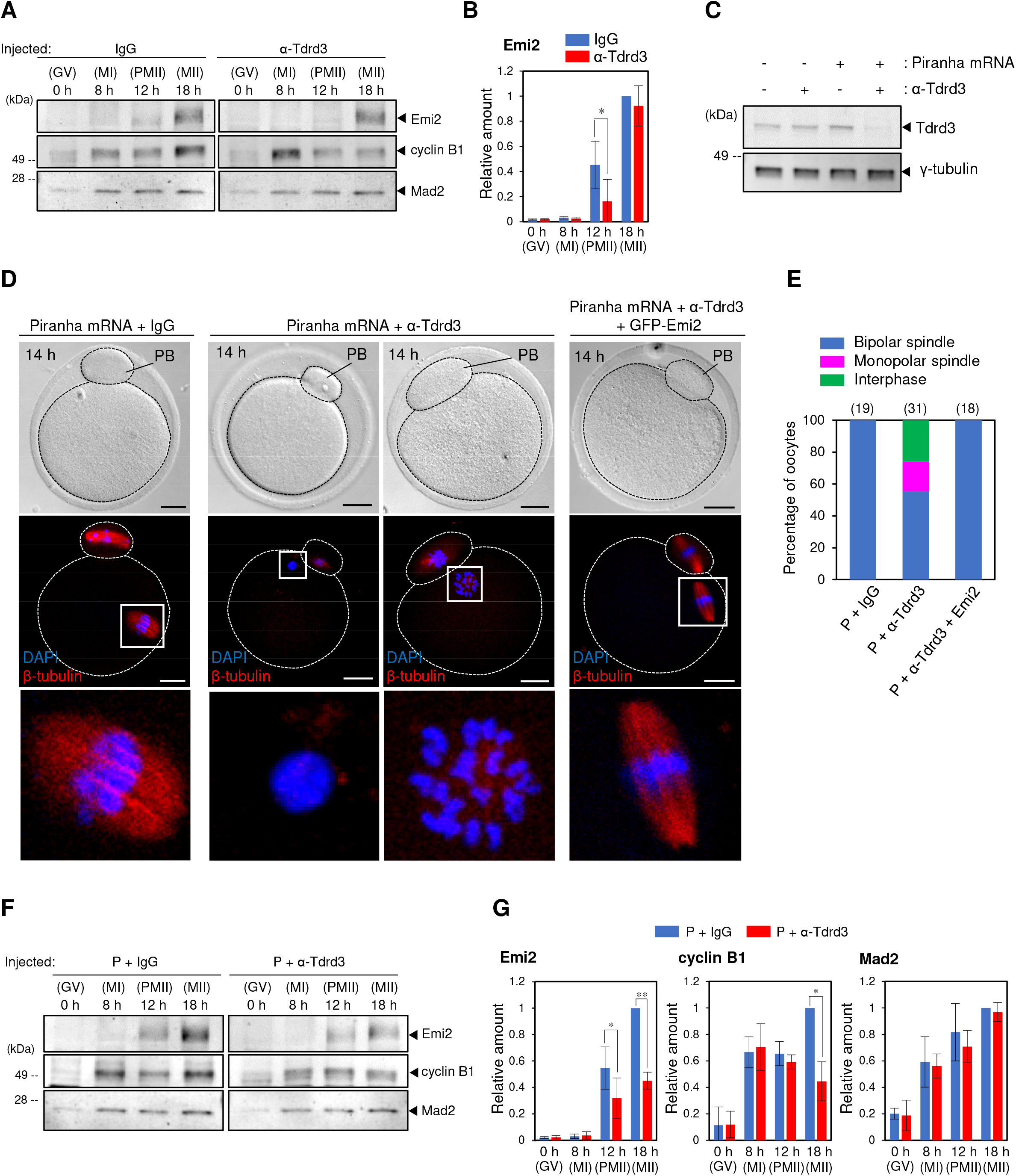
Knockdown of Tdrd3 attenuates Emi2 synthesis in meiosis II but not cyclin B1 and Mad2 synthesis in meiosis I. (A) Immunoblotting of Emi2, cyclin B1 and Mad2 in oocytes injected with IgG or anti-Tdrd3 antibody at 0, 8, 12, and 18 h after resumption of meiosis. (B) Quantitative analysis for the amount of Emi2 in (A) (means ± SD; n =3). *t*-test: * *P* < 0.05. (C) Immunoblotting of Tdrd3 and γ-tubulin in oocytes not injected (-) and injected (+) with Piranha mRNA and anti-Tdrd3 antibody at 18 after resumption of meiosis. (D) Immunofluorescence of β-tubulin in oocytes injected with Piranha mRNA and IgG (left), Piranha mRNA and anti-Tdrd3 antibody (middle), or Piranha mRNA, anti-Tdrd3 antibody and GFP-Emi2 mRNA at 14 h after resumption of meiosis. Lower panels show enlarged views of the boxed region. DNA is shown in blue. PB, polar body. Bars: 20 μm. (E) Percentage of oocytes with a bipolar or monopolar spindle or those that entered interphase in oocytes injected with Piranha mRNA and IgG (P + IgG), Piranha mRNA and anti-Tdrd antibody (P + α-Tdrd3), or Piranha mRNA, anti-Tdrd3 antibody and GFP-Emi2 mRNA (P + α-Tdrd3 + Emi2) at 14 h after resumption of meiosis. (F) Immunoblotting of Emi2, cyclin B1 and Mad2 in oocytes injected with Piranha mRNA and IgG (P + IgG) or Piranha mRNA and anti-Tdrd3 antibody (P + α-Tdrd3) at 0, 8, 12, and 18 h after resumption of meiosis. (G) Quantitative analyses for the amounts of Emi2, cyclin B1 and Mad2 in (F) (means ± SD; n =3). *t*-test: * *P* < 0.05, ** *P* < 0.01.

Injection of mRNA encoding a modified version of TRIM21, Piranha, with the anti-Tdrd3 antibody effectively reduced the amount of Tdrd3, while no reduction was observed with injection of *Piranha* mRNA with control IgG (Fig. 6C). These oocytes formed a pronucleus-like structure and a monopolar spindle after extruding the polar body, while no defect was observed in control oocytes injected with *Piranha* mRNA and IgG (Fig. 6D and E). The accumulation of Emi2 was attenuated throughout meiosis II by the knockdown of Tdrd3 (Fig. 6F and G). In contrast, the amounts of cyclin B1 and Mad2 in meiosis I were not affected (Fig. 6F and G). The amount of cyclin B1, but not that of Mad2, was reduced in meiosis II, which would be caused by the activation of APC/C in meiosis II due to the reduced levels of Emi2. The defect in meiosis II was almost completely recovered by GFP-Emi2 expressed after completion of meiosis I (Fig. 6D and E). Taken together, these results indicate that Tdrd3 is required for the translation of *Emi2* mRNA in meiosis II and regulates the progression of meiosis II thorough the synthesis of Emi2 protein after completion of meiosis I until MII arrest.

## DISCUSSION

In *Xenopus* oocytes, Emi2 protein has been shown to accumulate after meiosis I and prevent APC/C during the progression of meiosis II (Liu et al., 2006; Ohe et al., 2007; Tung et al., 2007). Analyses using reporter mRNAs suggested that the translational control of *Emi2* mRNA, which would be dependent on CPE and CPEB1, was important for the timing of Emi2 accumulation (Ohe et al., 2007; Tung et al., 2007). However, whether the translation of endogenous *Emi2* mRNA is involved in the timely controlled protein accumulation has been obscure. Moreover, whether *Emi2* mRNA is regulated at the translational level in mouse oocytes has remained completely unknown. In this study, we demonstrated that *Emi2* mRNA is translationally repressed with short poly(A) tails in meiosis I and translationally activated in a polyadenylation-dependent manner in meiosis II in mouse oocytes (Fig. 1). Tdrd3 was identified as a novel protein binding specifically to *Emi2* mRNA and was shown to colocalize with *Emi2* RNA granules in GV-stage oocytes (Figs. 2, 3, 4 and 5). Knockdown of Tdrd3 caused failure in meiosis II entry and MII spindle formation through attenuation of Emi2 synthesis (Fig. 6). Collectively, our results provide evidence that the Emi2 accumulation in meiosis II is actually regulated by the translational control of *Emi2* mRNA that is deposited in oocytes. Moreover, Tdrd3-mediated translation of *Emi2* mRNA is fundamental for the generation of a fertilizable gamete.

FISH analyses of mouse ovaries showed that *Emi2* mRNA assembled into RNA granules, which are different from those of cyclin B1, in GV-stage oocytes (Fig. 2). IP/RT-PCR and the RNA pull-down assay followed by mass spectrometry analysis suggested that *Emi2* and cyclin B1 RNA granules share certain proteins, but each granule also possesses specific components (Figs. 3 and 4). These results suggest that accurate translational control of assembled mRNAs is achieved by the assembly of common and specific components of RNA granules. In contrast to the specific components such as Pum1 and Tdrd3, HuR and HuB were shown to bind to both cyclin B1 and *Emi2* mRNAs (Fig. 3B and C). HuR is widely expressed in various cells and tissues and has been shown to be a component of stress granules (Gallouzi et al., 2000; Papadopoulou et al., 2013). In HuR-knockdown cultured human cells, the size of stress granules generally became much smaller, and the number of granules decreased, indicating that HuR is a key component in assembling stress granules (Fujimura et al., 2009). HuB is expressed specifically in neurons and in the testis and ovary. Recent studies have shown that HuB is a component of neuronal stress granules (Markmiller et al., 2018) and functions in translational repression of many mRNAs in mouse oocytes (Chalupnikova et al., 2014). Therefore, HuR and HuB may function in assembling and translationally repressing cyclin B1 and *Emi2* mRNAs in mouse oocytes. Our UV cross-linking assay showed that CPEB1 also binds to both *Emi2* and cyclin B1 mRNAs (Fig. 4C). In cultured human cells, CPEB1 was shown to be assembled in stress granules (Wilczynska et al., 2005). In contrast to Pum1 and Tdrd3, HuR, HuB and CPEB1 would be constitutional components of RNA granules in mouse oocytes regardless of the timing of translational activation of assembled mRNAs.

The *tudor* gene was identified in *Drosophila* as one of the maternal factors that regulate embryonic development and fertility (Boswell and Mahowald, 1985). The tudor domain forms a β-barrel-like core structure that contains three short β-strands followed by an a-helical region, and over 200 tudor domain-containing proteins have been identified from many eukaryotes including fungi, plants and animals (Lasko, 2010; Ponting, 1997). Many tudor domain-containing proteins harbor RNA-binding motifs, chromatin-binding domains, or DNA-binding domains and have been shown to function in diverse cellular processes such as histone modification, DNA damage response, RNA splicing, small RNA pathway, and pi-RNA-mediated transposon silencing (Maurer-Stroh et al., 2003; Siomi et al., 2010). The full length of Tdrd3 was identified on the basis of its tudor domain and has been shown to possess several domains, which may be involved in interaction with nucleic acid and/or proteins (Goulet et al., 2008; Linder et al., 2008). Using cultured human cells, Tdrd3 was shown to assemble into stress granules in response to several types of stress. In breast cancer cells, Tdrd3 was shown to selectively regulate the translation of certain mRNAs, which may promote growth and invasion of breast cancer tumors (Morettin et al., 2017). However, whether Tdrd3 functions in physiological conditions has remained unknown.

In this study, we identified Tdrd3 as a protein binding specifically to *Emi2* mRNA by the RNA pull-down assay followed by mass spectrometry analysis (Fig. 3D and Table S1). We showed that the N-terminal region of Tdrd3 (1-473 aa) bound to the 3’UTR of *Emi2* mRNA in *in vitro* UV cross-linking analysis (Fig. 4D), indicating that this region contains the amino acid sequences responsible for the direct binding to mRNA. In addition, this region is able to select the target mRNA because Tdrd3-N did not bind to the 3’UTR of cyclin B1 mRNA (Fig. 4D). Our results provide the first evidence that Tdrd3 directly binds to mRNA and has target specificity. In GV-stage mouse oocytes, Tdrd3 formed particles and colocalized with *Emi2* RNA granules (Fig. 5). Interference in Tdrd3 function by the anti-Tdrd3 antibody delayed but did not completely inhibit Emi2 synthesis in meiosis II (Fig. 6). Knockdown of Tdrd3 did not induce or accelerate the derepression of *Emi2* mRNA translation (Fig. 6), suggesting that Tdrd3 is not a repressor of translation. Rather, it seems to promote the translation of *Emi2* mRNA in meiosis II, since knockdown attenuated the synthesis of Emi2 in meiosis II. Intriguingly, both interference in Tdrd3 function by the anti-Tdrd3 antibody and knockdown of Tdrd3 did not affect the translation of cyclin B1 and *Mad2* mRNA in meiosis I (Fig. 6). These results are consistent with the results of the IP/RT-PCR, RNA pull-down assay, UV cross-linking, and FISH analyses (Figs. 3, 4 and 5) and support the notion that the accurate timing of translational activation of mRNA is determined by components specific to distinct RNA granules.

Super-resolution microscopy of Tdrd3 and Pum1 in GV-stage mouse oocytes showed differences in the sizes and shapes of Tdrd3 particles and Pum1 aggregates (Fig. 5). Pum1 was shown to form solid-like aggregates in human cultured cells and GV-stage mouse oocytes (Shiina, 2019; Takei et al., 2020). We previously showed that highly structured aggregates of Pum1 in a solid-like state ensures translational repression of cyclin B1 and *Mad2* mRNAs and that dissolution of the aggregates allows the disassembly of cyclin B1 and *Mad2* RNA granules, resulting in the translational activation of these mRNAs (Takei et al., 2020). The quantitative analyses showed that the size of Tdrd3 particles was smaller than that of Pum1 aggregates (Fig. 5). Moreover, Tdrd3 particles appeared in a round shape (Fig. 5), which might reflect the formation of liquid droplets induced by liquid-liquid separation (Shin and Brangwynne, 2017). Thereby, Tdrd3 might form soluble particles and act in a way different from that of Pum1. We are currently addressing this possibility. Interestingly, Tdrd3 particles seemed to form clusters in the oocyte cytoplasm (Figs. 5 and S3) and these distribution patterns were maintained in meiosis I (T. Kotani, unpublished observations/data not shown). The highly organized complexes including Tdrd3 might promote compartmentalization of the cytoplasm and maintain silencing of translation in meiosis I. After completion of meiosis I, Tdrd3 promotes the *Emi2* mRNA translation by an unknown mechanism. One possible explanation is that Tdrd3 recruits Dazl and/or poly(A) polymerases such as mGLD-2 after completion of meiosis I, both of which have been shown to accumulate in meiosis I and to promote translation of many dormant mRNAs in mouse oocytes (Chen et al., 2011; Nakanishi et al., 2006; Sousa Martins et al., 2016).

In conclusion, we demonstrated the existence of Tdrd3-mediated translational control of *Emi2* mRNA, through which oocytes promote the progression of meiosis II, generating fertilizable oocytes. The highly ordered temporal translation of mRNAs would be achieved by the assembly of components common and specific to distinct RNA granules. These granules were spatially and temporally organized in the oocyte cytoplasm. Recent studies utilizing systematic approaches have demonstrated that more than one hundred proteins are assembled in stress granules and that stress granules compose diversities; specific proteins in addition to constitutive proteins are assembled by different types of stress (Aulas et al., 2017; Jain et al., 2016; Markmiller et al., 2018). Compositional differences in stress granules were also observed in different types of cells such as fibroblasts and neuronal cells. Our results will contribute to an understanding of the differences in function and regulation of these granules in many types of cells besides oocytes.

## MATERIALS AND METHODS

### Preparation of ovaries

All animal experiments in this study were approved by the Committee on Animal Experimentation, Hokkaido University. Mouse ovaries were dissected from 8- to 12-week-old adult females and placed in PBS (137 mM NaCl, 2.7 mM KCl, 10 mM Na2HPO4, and 2 mM KH2PO4, pH 7.2). For *in situ* hybridization, mouse ovaries were fixed with 4% paraformaldehyde in PBS (4% PFA/PBS) overnight at 4°C. For immunoprecipitation analysis, mouse ovaries were homogenized with an equal volume of ice-cold extraction buffer (EB: 100 mM β-glycerophosphate, 20 mM Hepes, 15 mM MgCl2, 5 mM EGTA, 1 mM dithiothreitol, 100 μM (p-amidinophenyl) methanesulfonyl fluoride, and 3 μg/ml leupeptin, pH 7.5) containing 1% Tween 20 and 100 U/ml RNase inhibitor (Invitrogen). After centrifugation at 5,000 rpm for 15 min at 4°C, the supernatant was collected and used for immunoprecipitation. For the RNA pull-down assay, cytoplasmic extracts of *Xenopus* oocytes were prepared according to the procedure of the RiboTrap kit (MBL International). Ovaries were dissected from adult females and cut into small pieces in nuclease-free PBS. *Xenopus* ovaries were homogenized with an equal volume of ice-cold CE Buffer (+) (MBL International) and incubated for 10 min on ice. Then the extracts were diluted with Detergent Solution (MBL International) and centrifuged at 3,000 g for 3 min at 4°C. The supernatant was transferred to new tubes and diluted with High-Salt Solution (MBL International). After centrifugation at 12,000 g for 3 min at 4°C, the supernatant was collected and used for the RNA pull-down assay.

### Production of antibodies

DNA sequences encoding a part of mouse Emi2 (residues 1-366) was amplified by PCR and ligated into pET21 vector to produce a histidine (His)-tagged protein. The recombinant protein was expressed in *E. coli* and purified by SDS-PAGE, followed by electroelution in Tris-glycine buffer without SDS. The purified protein was dialyzed against 1 mM HEPES (pH 7.5), lyophilized, and used for injection into two mice. The obtained antisera were affinity-purified with recombinant Emi2-His protein electroblotted onto a membrane (Immobilon; EMD Millipore).

### Collection of oocytes

For immunoblotting, PAT assay and injection experiments, GV-stage oocytes were collected by puncturing ovaries with a needle in M2 medium (94.7 mM NaCl, 4.8 mM KCl, 1.2 mM KH2PO4, 1.3 mM MgCl2, 5.6 mM glucose, 23.3 mM sodium lactate, 4 mM NaHCO3, 0.3 mM sodium pyruvate, 1.7 mM CaCl2, 21 mM HEPES, 0.1 mM gentamicin, 4 mg/ml BSA, pH 7.5) supplemented with 10 μM milrinone (M2+). Resumption of meiosis was induced by washing oocytes with M2 medium lacking milrinone (M2-) four times and culturing at 37°C in drops of M2-covered with paraffin liquid. At the appropriate time points after resumption of meiosis, oocytes were collected for immunoblotting and the poly(A) test (PAT) assay. For immunoblot analysis, the collected oocytes were washed with PBS and extracted in 10 μl lithium dodecyl sulfate (LDS) sample buffer (NOVEX) or SDS sample buffer (50 mM Tris-HCl; pH 6.8, 2% sodium dodecyl sulfate, 6% β-mercaptoethanol, 10% glycerol, 0.1% phenol Red). For the PAT assay, oocytes were extracted with Trizol regent (Life Technologies) according to the manufacturer’s instructions.

### PAT assay

Two μg of total RNA extracted from pools of 200 mouse oocytes was ligated to 0.4 μg of P1 anchor primer (5’-P-GGT CAC CTT GAT CTG AAG C-NH2-3’) in a 10-μl reaction using T4 RNA ligase (New England Biolab) for 30 min at 37°C. The ligase was inactivated for 5 min at 92°C. Eight μl of the reaction was used in a 20 μl reverse transcription reaction using the SuperScript III First Strand Synthesis System with P1’ primer (5’-GCT TCA GAT CAA GGT GAC CTT TTT-3’). Aliquots of the cDNA were used as templates of the 1st RCR with primer sets of P1’ primer and m*Emi2*-forward primer-1 (5’-ATG TCT GTG CGC TTATCA CG-3’) or P1’ primer and mcyclin B1*-* forward primer-1 (5’-CCA CTC CTG TCT TGT AAT GC-3’). Aliquots of the 1st PCR reactions were used for the 2nd PCR with primer sets of P1’ primer and m*Emi2-*forward primer-2 (5’-TCC CAT TGA TGA CAG ACG TTG ATT TAC TTC CCA CTT-3’) or P1’ primer and mcyclin B1-forward primer-2 (5’-CCT GGA AAA GAA TCC TGT CTC-3’). The PCR products were resolved on 3% TAE gel. It was confirmed by cloning the 2nd PCR products and sequencing them that the increase in PCR product length was due to elongation of the poly(A) tails.

### Polysomal fractionation

Polysomal fractionation was performed according to the procedure described previously (Masek et al., 2020; Takada et al., 2020). Briefly, 200 oocytes were treated with 100 μg/ml of cycloheximide (CHX) for 10 min and collected in 350 μl of lysis buffer (10 mM Hepes, pH 7.5; 62.5 mM KCl, 5 mM MgCl2, 2 mM DTT, 1% TritonX-100) containing 100 μg/ml of CHX and 20 U/ml of Ribolock (Thermo Fisher Scientific). After disruption of the zona pellucida with 250 μl of zirconia-silica beads (BioSpec), lysates were centrifuged at 8,000 g for 5 min at 4°C. Supernatants were loaded onto 10-50% linear sucrose gradients containing 10 mM Hepes, pH 7.5; 100 mM KCl, 5 mM MgCl2, 2 mM DTT, 100 μg/ml of CHX, Complete-EDTA-free Protease Inhibitor (1 tablet/100 ml: Roche) and 5 U/ml of Ribolock. Centrifugation was performed using an Optima L-90 ultracentrifuge (Beckman) at 35,000 g for 65 min at 4°C. Polysome profiles were recorded using an ISCO UA-5 UV absorbance reader. Ten equal fractions were collected from each sample and subjected to RNA isolation by Trizol reagent (Sigma). Fractions 6 to 10 were taken for polysome-bound RNA. Furthermore, a library was prepared using the SMART-seq v4 ultra low input RNA kit (Takara Bio). Sequencing was performed by HiSeq 2500 (Illumina) as 150-bp paired-end. Reads were trimmed using Trim Galore v0.4.1 and mapped to the mouse GRCm38 genome assembly using Hisat2 v2.0.5. Gene expression was quantified as fragments per kilobase per million (FPKM) values in Seqmonk v1.40.0.

### Injection of MOs, mRNAs and antibodies

The sequences of antisense morpholino oligonucleotides (MOs) (Gene Tools, LLC) are as follows: *Emi2*-3’UTR-MO, 5’-ATTTATTTCTTAAAAGTCCCAACAA-3’, that specifically targets the terminal sequence of the *Emi2* mRNA 3’UTR and Emi2-3’UTR 5mm-MO, 5’-ATTCATTACTTAATAGTCACATCAA-3’, that contains 5-nts mismatches (underlines) and was used as a control. GV-stage oocytes were injected with 10 pl of a solution containing 0.1 mM Emi2-3’UTR-MO or Emi2-3’UTR 5mm-MO using an IM-9B microinjector (Narishige) under a Dmi8 microscope (Leica) in M2 medium. After being injected, the oocytes were cultured in M16 medium in an atmosphere of 5% CO2 in air at 37°C and used for immunoblotting or morphological observation after fixation with 4% PFA for 1 h.

mRNAs encoding GFP and GFP-Emi2 were synthesized using the mMESSAGE mMACHINE SP6 kit (Ambion) and dissolved in nuclease-free distilled water. Eight pl of 1 μg/μl mRNAs was injected into GV-stage oocytes in M2+ using the IM-9B microinjector under the Dmi8 microscope and washed four times with M2- to resume meiosis. Similarly, 8 pl of 0.1 mg/ml anti-Tdrd3 antibody (Cell Signaling Technology; D302G) or control IgG was injected into GV-stage oocytes and washed four times with M2-. To knock down Tdrd3 by the Trim-Away system, 8 pl of a mixture containing 500 ng/μl RFP-Piranha mRNA (System Biosciences) and 0.1 mg/ml anti-Tdrd3 antibody (Cell Signaling Technology; D302G) was injected into GV-stage oocytes and incubated at 37°C in M2+ for 2-3 h to allow Piranha protein expression. The oocytes were then washed four times with M2- to resume meiosis.

### Whole mount immunofluorescence

Oocytes were collected at the appropriate time points after incubating with M2- at 37°C. After fixation with 4% PFA in PBS for 1 h, the oocytes were permeabilized in permeabilization buffer (0.3% BSA, 0.1% Triton-X, 0.02% NaN3 in PBS) for 20 min, washed three times with blocking solution (0.3% BSA, 0.01% Tween 20 in PBS), and incubated in blocking solution for 20 min at room temperature. The oocytes were then incubated with anti-β-tubulin-Cy3 antibody (Sigma) for 1 h at room temperature. After washing three times with blocking solution, the oocytes were mounted on slides with VECTASHIELD Mounting Medium containing DAPI (Vector Laboratories, lnc) and observed under an LSM 5 LIVE confocal microscope (Carl Zeiss).

### Section *in situ* hybridization

Section *in situ* hybridization with the TSA Plus system (PerkinElmer) was performed according to the procedure reported previously (Takei et al., 2018). In brief, the fixed ovaries were dehydrated, embedded in paraffin, and cut into 7-μm-thick sections. The digoxigenin (DIG)-labeled sense or antisense RNA probe for full-length *Emi2* was synthesized with a DIG-RNA-labeling kit (Roche Molecular Biochemicals). No signal was detected with sense probes. The sections of mouse ovaries were hybridized with a hybridization mix containing 1 ng/μl of DIG-labeled RNA probe. After hybridization and washing, the sections were incubated with anti-DIG-horseradish peroxidase (HRP) antibody (1:500 dilution; Roche Molecular Biochemicals) for 30 min. The reaction with tyramide-DNP, followed by reaction with the anti-DNP-alkaline phosphatase (AP) antibody, and the detection of signals were performed according to the manufacturer’s instructions. For FISH analysis, the samples were incubated overnight with anti-DNP-Alexa Fluor 488 antibody (1:500 dilution; Molecular Probes) after reaction with tyramide DNP. To detect nuclei, the samples were incubated with 10 μg/ml Hoechst 33258 for 10 min.

Double fluorescence *in situ* hybridization of cyclin B1 and *Emi2* mRNAs was performed as follows. Sections of mouse ovaries were hybridized with a fluorescein-labeled cyclin B1 RNA probe and DIG-labeled *Emi2* RNA probe. After hybridization and washing, samples were incubated with anti-DIG-HRP antibody (1:500 dilution) for 30 min. After reaction with tyramide-DNP, the samples were incubated with anti-DNP-Alexa Fluor 488 antibody (1:500 dilution) overnight. Before detection of the fluorescein-labeled cyclin B1 RNA probe, the samples were incubated with 1% H2O2 in PBS for 15 min for inactivating HRP. The reaction with tyramide-Cy3 (1:50 dilution in 1 ×Plus Amplification Diluent (PerkinElmer), followed by 1:100 dilution with DW) was performed for 20 min. To detect nuclei, the sections were incubated with 10 μg/ml Hoechst 33258 for 10 min and observed under the LSM 5 LIVE confocal microscope.

### Section immunofluorescence

Section immunofluorescence was performed according to the procedure reported previously (Takei et al., 2020). Briefly, ovary sections were microwaved for 10 min (500 W) with 10 mM citric acid (pH 6.0). After incubation with a TNB blocking solution (PerkinElmer) at room temperature for 1 h, the samples were incubated with anti-TDRD3 antibody (Cell Signaling Technology; D302G) and/or anti-Pum1 (Bethyl; A300-201A) antibody overnight at room temperature. The samples were then incubated with anti-goat IgG-Alexa Flour Plus 555 antibody (Invitrogen) and/or anti-goat IgG-Alexa Flour Plus 647 antibody (Invitrogen) at room temperature for 1 h. After staining with Hoechst 33258, the samples were mounted and observed under an N-SIM super-resolution microscope (Nikon). No signal was detected in the reaction without primary antibodies. To simultaneously detect the protein and mRNA, the samples were immunostained as described above after detection of the *Emi2* or cyclin B1 RNA probe with tyramide-FITC (PerkinElmer, Inc.) in *in situ* hybridization analysis. The size and roundness of signals were measured by ImageJ software. The creation of random dots and measurements of the distances of Tdrd3 particles and randomly distributed dots were performed by using ImageJ.

### Immunoprecipitation followed by RT-PCR (IP/RT-PCR)

Eighty μl of mouse ovary extracts was incubated with 2 μg of anti-Pum1 (Bethyl; A300-201A), anti-HuR (SANTA CRUZ; sc-5261) or anti-ELAVL2 (HuB) (Proteintech; 14008-1-AP) antibodies and protein G Mag sepharose (GE Healthcare) overnight at 4°C. the same volume of IgG was used as a control. The samples were washed five times with EB containing 1% Tween 20. An aliquot of the samples was used for RT-PCR with primer sets specific to cyclin B1, mcyclin B1*-* forward primer-2 (5’-CCT GGA AAA GAA TCC TGT CTC-3’) and mcyclin B1*-* reverse primer-1, specific to *Emi2*, m*Emi2-*forward primer-2 (5’-TCC CAT TGA TGA CAG ACG TTG ATT TAC TTC CCA CTT-3’) and *Emi2*-reverse primer (5’-GGG GAT ATG ACA TAG TAA AA-3’), and specific to *α-tubulin, ma-tubulin-forward* primer (5’-CTT TGT GCA CTG GTA TGT GGG T-3’) and *ma-tubulin-reverse* primer (5’-ATA AGT GAA ATG GGC AGC TTG GGT-3’).

### RNA pull-down assay and mass spectrometry

The RNA-binding assay was performed using the RiboTrap Kit (MBL International) according to the manufacturer’s instructions. In brief, bromouridine (BrU)-labeled RNAs of the 3’ UTR of mouse *Emi2, Xenopus Emi2*, and mouse cyclin B1 were generated using the Riboprobe *in vitro* Transcription Systems kit according to the manufacturer’s protocol (Promega). Anti-BrdU antibodies were conjugated with protein A-sepharose beads overnight at 4°C (GE Healthcare). Then the RNAs were bound to the beads. Cytoplasmic extracts of *Xenopus* oocytes were transferred to tubes containing the BrU-labeled RNAs conjugated with the beads for 2 h at 4°C. The samples were washed with Wash BufferII, eluted with elution buffer (4% BrdU/DMSO solution in nuclease-free PBS; MBL International), and subjected to SDS-PAGE. The SDS-PAGE gels were stained by silver staining with the Silver Stain MS kit (Wako Pure Chemical Industries, Ltd) to visualize the proteins associated with BrU-labeled RNAs. The bands were excised from the gel and subjected to a mass spectrometric analysis with Orbitrap Velos Pro (Thermo Fisher Scientific). For protein identification, MASCOT 2.5.1 was used for database searching against NCBInr_*Xenopus laevis* (updated on 05/09/2015, 17,456 sequences). The results from each run were filtered with the peptide confidence value, in which peptides showing a false discovery rate (FDR) of less than 1% were selected. In addition, proteins identical to the sequenced peptides with the rank1 values were selected. The number of peptides identical to the proteins was more than one. We counted the number of proteins that were pulled down with the mouse and *Xenopus Emi2* RNA probes as candidate proteins specifically binding to *Emi2* mRNA, with all of the RNA probes as candidates binding to both *Emi2* and cyclin B1 mRNAs, and with the mouse cyclin B1 RNA probe as candidates specifically binding to cyclin B1 mRNA.

### UV-cross linking assay

The full length of Tdrd3 was amplified by PCR using Tdrd3-f1 (5’-TGG ATC CTA TGG CCG AGG TGT CCG-3’) and Tdrd3-r1(5’-ATC GAT GTT CCG AGC TCG AGG-3’). The N-terminal region of Tdrd3 (1-473 aa) was amplified by PCR using Tdrd3-f1 and Tdrd3-r2 (5’-ATC GAT TGG CCT GTC ACA CTT-3’). The C-terminal region of Tdrd3 (246-744 aa) was amplified by PCR using Tdrd3-f2 (5’-GGA TCC TAT GGG TGG TGC CAG AAG TAA-3’) and Tdrd3-r1. The PCR products were inserted into pGEM-T Easy vector (Promega). Sequences encoding the full length and parts of Tdrd3 were cloned into pCS2-Flag-N or pCS2-Flag-C to produce Tdrd3 fused with Flag at the N- or C-terminus of Tdrd3. mRNAs encoding Flag-Tdrd3, Flag-Tdrd3-N and Flag-Tdrd3-C were synthesized with an mMESSAGE mMACHINE SP6 kit (Ambion), and the resulting mRNAs (2 μg) were translated in 50 μl of rabbit reticulocyte lysate (Promega). Flag-tagged proteins were purified by immunoprecipitation with anti-Flag M2 antibody (Sigma) and Protein G Mag Sepharose (GE healthcare). The DIG-labeled 3’UTR of *Emi2* and cyclin B1 RNA probes were synthesized with a DIG-RNA-labeling kit (Roche Molecular Biochemicals). Two μg of probes was incubated with purified Flag-tagged proteins for 15 min, and the reaction mixtures were irradiated twice in a UV cross-linker (XL-1000, Funakoshi) at an energy setting of 86 mJ/cm^2^.

### Immunoblotting

Mouse oocyte extracts were separated by SDS-PAGE with Bolt Bis-Tris Plus Gels (Novex), blotted onto an immobilon membrane using a Bolt Mini Blot Module (Novex), and probed with anti-Emi2 (this study), anti-cyclin B1 (Abcam; V152), anti-γ-tubulin (Sigma; T6557), anti-Tdrd3 (Cell Signaling Technology; D302G) and anti-Mad2 (Bethyl; A300-301A) antibodies. The crude extracts from mouse ovaries and rabbit reticulocyte lysates and the immunoprecipitates were separated by SDS-PAGE, blotted onto an immobilon membrane, and probed with anti-Pum1 (Bethyl; A300-201A), anti-HuR (SANTA CRUZ; sc-5261), anti-ELAVL2 (HuB) (Proteintech; 14008-1-AP), anti-FLAG (Sigma; F1804-1MG), anti-TDRD3 (Cell Signaling Technology; D302G) and anti-DIG-AP (Roche) antibodies. The intensity of signals was quantified using ImageJ software.

## Acknowledgements

We thank Dr. K. Kobayashi for technical advice on super-resolution microscopy and S. Kawamura for an initial experiment of the PAT assay.

## Author contributions

Conceptualization: N. Takei and T. Kotani. Investigation: N. Takei, K. Sato, Y, Takada and R. Iyyappan. Resources: A. Susor and T. Yamamoto. Project administration: T. Kotani. Writing - original draft: N. Takei and T. Kotani. Writing - review and editing: A. Susor, T. Yamamoto and T. Kotani.

